# “Continuous Neural Correlates of Imbalanced Reinforcement Learning in Obsessive-Compulsive Disorder and Healthy Individuals”

**DOI:** 10.1101/2025.01.28.635211

**Authors:** Yuki Sakai, Yutaka Sakai, Yoshinari Abe, Jin Narumoto, Saori C. Tanaka

**Affiliations:** ATR Brain Information Communication Research Laboratory Group, 2-2-2 Hikaridai Seika-Cho, Soraku-Gun, Kyoto 619-0288, Japan; Department of Psychiatry, Graduate School of Medical Science, Kyoto Prefectural University of Medicine, 465 Kajii-Cho, Kawaramachi-Hirokoji, Kamigyo-Ku, Kyoto 602-8566, Japan; Brain Science Institute, Tamagawa University, 6-1-1, Tamagawa-gakuen, Machida, Tokyo 194-8610, Japan; Division of Information Science, Graduate School of Science and Technology, Nara Institute of Science and Technology, 8916-5 Takayama-Cho, Ikoma, Nara 630-0192, Japan

**Keywords:** Computational model, Functional Connectivity, Functional Magnetic Resonance Imaging, Obsessive-Compulsive Disorder (OCD), Reinforcement Learning

## Abstract

**Aim:** Obsessive-compulsive disorder (OCD) is characterized by imbalanced reinforcement learning. This study investigated neural correlates of this imbalance by examining resting-state functional connectivity (rs-fMRI) in non-medicated OCD patients and healthy controls (HCs) exhibiting varying degrees of memory trace imbalance.

**Methods:** We employed network-based statistics (NBS), suitable for identifying network-level alterations of rs-fMRI data from 49 OCD patients and 53 HCs (Core Discovery Dataset: Core-DS) to identify a significant network. We validated this network in an independent dataset of 10 OCD patients and 18 HCs (Independent Validation Dataset: IndV-DS). Additionally, we compared functional connectivity in the identified network between subgroups of HCs with imbalanced (OCD-like) and balanced learning profiles (Extension Dataset: Ext-DS; n=10 each).

**Results:** NBS identified an ‘OCD network’ with increased connectivity in OCD patients (p_adjusted_ = 0.022), comprising regions in the dorsolateral prefrontal cortex (DLPFC), parietal cortex, retrosplenial cortex, and hippocampal formation. This network’s increased connectivity in OCD patients was replicated in IndV-DS (p = 0.0027). Importantly, imbalanced HCs showed a significant increase in functional connectivity, particularly between the DLPFC and presubiculum, in the identified ‘OCD network’, compared to balanced HCs (p = 0.0051, p_adjusted_ = 0.066).

**Conclusion:** Increased functional connectivity in the ‘OCD network,’ especially between the DLPFC and presubiculum, is present in both OCD patients and a subgroup of HCs with learning imbalances. These findings provide evidence for a continuous neural basis of imbalanced reinforcement learning, suggesting a continuum of traits associated with OCD between patients and “healthy” individuals.

## Introduction

Obsessive-compulsive disorder (OCD) is characterized by persistent, unwanted, intrusive thoughts (obsessions) that trigger repetitive behaviors or mental acts (compulsions) aimed at reducing distress. This disorder presents a significant challenge to public health and is a substantial burden, as even subthreshold symptoms pose a considerable challenge ^1^. Resting-state functional magnetic resonance imaging (rs-fMRI) has become a valuable tool for investigating neural underpinnings of psychiatric disorders, offering the possibility of identifying clinically relevant biomarkers and probing functional connectivity (FC) alterations ^2-4^. Previous rs-fMRI studies have reported widespread functional dysconnectivity in OCD involving the frontoparietal and salience networks and altered connectivity between salience, frontoparietal, and default mode networks ^5^. Independently, computational models have begun to explore mechanisms of OCD, proposing that imbalances in reinforcement and punishment systems may contribute to development and maintenance of OCD ^6^.

Despite these advances, several critical questions remain regarding precise neural mechanisms underlying OCD. While rs-fMRI studies have identified altered FC in OCD, the link between these network-level findings and specific computational processes, such as those related to reinforcement learning, remains unclear. Specifically, our previous computational modeling work demonstrated that an imbalance in reinforcement and punishment learning, indexed by trace decay factors, could reinforce implicit choices, potentially leading to OCD-like behaviors ^6^. Moreover, we found that this imbalance, indexed by trace decay factors, exists on a continuum among OCD patients and healthy controls (HCs), with HCs showing imbalanced learning parameters exhibiting significantly stronger obsessive-compulsive tendencies than those with balanced parameters. These findings and our prior study suggest a behavioral continuum, but whether a continuous neural basis supports this remains unknown. Thus, a critical gap exists in understanding whether a continuous neural basis underpins the behavioral imbalance seen among both OCD and a subgroup of HCs and whether this neural basis consists of abnormalities typically associated with OCD. While other computational models of OCD exist ^7-9^, our model uniquely explains the cyclical relationship between obsessions and compulsions and understands mechanisms of action of behavioral therapy and pharmacological interventions while also capturing the continuity with subclinical obsessive-compulsive traits found in the general population. Addressing this gap, specifically, whether a neural continuum exists that mirrors the behavioral continuum, is crucial, as it can help clarify the complex pathophysiology of OCD and provide a more nuanced understanding of the obsessive-compulsive spectrum. Such insights can potentially inform development of more targeted therapeutic and preventative interventions.

To understand neural mechanisms of these behavioral imbalances, reinforcement learning principles, particularly credit assignment—linking actions to outcomes ^10^, are crucial. Although regions such as the dorsolateral prefrontal cortex (DLPFC) and retrosplenial cortex (RSC), thought to be involved in credit assignment and memory retrieval, are crucial for reinforcement learning ^11-13^, how these regions contribute to imbalanced learning remains unclear. Therefore, using a data-driven approach, we examined whole-brain, resting-state, FC in OCD and HC to identify neural substrates of OCD and imbalanced reinforcement learning.

## Methods

### Participants

This study included three independent datasets. Core Discovery Dataset (Core-DS) comprised 49 non-medicated OCD patients and 53 healthy controls (HCs) recruited at the Kajiicho Medical Imaging Center. Independent Validation Dataset (IndV-DS), used for replication, included 10 non-medicated OCD patients and 18 HCs recruited at the Kyoto Prefectural University of Medicine (KPUM). Extension Dataset (Ext-DS) comprised 20 HCs who had previously completed a delayed feedback task ^6^; based on their learning profiles, these participants were divided into two groups: 10 with imbalanced learning (*ν*^+^ > *ν*^−^) and 10 with balanced learning. This grouping was based on data-driven clustering in a larger dataset and has not changed from our previous study ^6^. Importantly, no participant was included in more than one dataset. All demographic distributions were matched between groups in each dataset, as detailed in Supplementary Tables 1, 2, and 3. Some participants in Core-DS, IndV-DS, and Ext-DS had previously participated in unrelated studies ^3, 6, 14-16^. A summary of fMRI imaging protocols using gradient EPI sequences for all datasets can be found in Supplementary Table 4, with high-resolution T1-weighted structural images also acquired for each participant. All participants provided written informed consent after receiving a complete study description. The protocol was approved by the Medical Committee on Human Studies at KPUM and the Ethics Committee at ATR. OCD diagnoses were primarily determined using the Structured Clinical Interview for DSM-IV Axis I Disorders-Patient Edition (SCID) ^17^, and symptom severity was assessed using the Yale-Brown Obsessive-Compulsive Scale (Y-BOCS) ^18^. Participants were excluded for metallic implants, significant medical conditions, prior psychosurgery, intellectual disability (IQ < 80), current primary psychiatric disorder diagnosis other than OCD, or pregnancy. Medication exclusion criteria were limited to psychotropic medications other than serotonin reuptake inhibitors (SRIs). Handedness was classified based on a modified 25-item version of the Edinburgh Handedness Inventory.

### Behavioral task and parameter estimation in Ext-DS

Data for the behavioral task, designed to probe the effect of reinforcement and punishment on delayed feedback learning, were acquired from a previous study ^6^. In this two-choice, delayed feedback task, participants chose between two abstract cues, each associated with varying magnitude (reward or punishment) and feedback delay.

The same reinforcement learning model as described in Sakai et al. ^6^ was used to fit each participant’s choice data individually. This model, detailed further in that publication, uses a separate eligibility trace for positive and negative prediction errors. Briefly, each trace represents the recent frequency of action in a specific state in a time scale determined by the trace decay factor, *ν*. If eligibility traces for positive and negative prediction errors are implemented in distinct neural systems, the respective eligibility traces 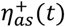 and 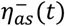 at time step *t* for action *a* (*a* = 0, 1, …) in state *s* (*s* = 0, 1, …) obey the following equation ^10^,

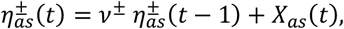

where *X*_*as*_(*t*) = 1 when action *a* is chosen in state *s* at time step *t* and *X*_*as*_ (*t*) = 0 otherwise. When a prediction error occurs, the choice probability of each action in each state is updated by the product of the eligibility trace and the prediction error. Thus, a positive prediction error reinforces recent actions in a time scale, whereas a negative error punishes them. Theoretically, trace factors in the two systems, *ν*^±^ , should be balanced: *ν*^+^ = *ν*^−^ . However, separate neural implementations make it difficult to maintain the balance perfectly. In our separate eligibility trace model, we assumed that an imbalance in trace factors could occur: *ν*^+^ ≠ *ν*^−^ . Model parameters were estimated using a maximum a posteriori method. Precise details of the task design, stimuli, and parameter estimation procedures can be found in the original publication ^6^.

### Preprocessing and statistical analysis of rs-fMRI data

Preprocessing steps, including slice timing correction, motion correction, and spatial normalization to the Montreal Neurological Institute (MNI) space, were performed using fmriprep ^19, 20^. Subsequently, data were converted from the volumetric NIfTI format to the surface-based CIFTI format (https://www.nitrc.org/projects/cifti/) using ciftify ^19^. To reduce artifacts and enhance the signal-to-noise ratio, rs-fMRI time series were detrended, bandpass filtered (0.01–0.08 Hz), and nuisance signals—including whole-brain signal fluctuations, six head motion parameters and their derivatives, and six anatomical CompCor components ^21^—were regressed out. With respect to motion artifacts, framewise displacement (FD) did not differ significantly between groups in any dataset [median (interquartile range); Core-DS: OCD patients, 0.082 (0.070–0.097) mm; HCs, 0.087 (0.067–0.10) mm; Brunner-Munzel test, p > 0.05; IndV-DS: OCD patients, 0.095 (0.067–0.12) mm; HCs, 0.079 (0.062–0.094) mm; Brunner-Munzel test, p > 0.05; Ext-DS: balanced subgroup, 0.13 (0.11–0.18) mm; imbalanced subgroup, 0.11 (0.094–0.14) mm; Brunner-Munzel test, p > 0.05]. The first six functional scans were discarded to allow magnetization to reach equilibrium. For each participant, mean time series were extracted from 360 cortical and 358 subcortical parcels using Cole-Anticevic Brain-wide Network Partition (CAB-NP) ^22^, which is a comprehensive whole-brain solution for large-scale functional networks based on the cortical parcellation developed by Glasser et al. ^23^ (Human Connectome Project Multi-Modal Parcellation). Pearson correlation coefficients were calculated between each pair of parcels and transformed to Fisher’s Z scores to obtain the FC matrix.

To evaluate the neural substrate of non-medicated patients with OCD hypothetically related to v^+^ > v^−^, we compared FC matrices between 49 non-medicated OCD patients and 53 HCs (Core-DS) using network-based statistics (NBS). We used NBS as they enable identification of interconnected subnetworks associated with the experimental condition while controlling the family-wise error rate ^24^. Briefly, NBS involved two steps: first, between-group comparisons were performed for all possible FCs and thresholded at t = 3.87 (p = 0.0001), controlling for FD. Thresholded subnetworks were then defined, and their size was calculated. Second, subnetwork significance was assessed via 10,000 random group permutations to estimate the null distribution of the largest subnetwork size. Subnetworks exceeding the family-wise error-corrected p-value of 0.05 were considered significant. The identified subnetwork was visualized using BrainNet Viewer ^25^. To validate the identified OCD network, mean FC in this network was compared between 10 OCD patients and 18 HCs from IndV-DS, using the Brunner-Munzel test. To determine if the imbalanced (*ν*^+^ > *ν*^−^) HC subgroup also showed altered FC in the identified OCD network, we compared all FCs in this network between 10 HCs in the imbalanced subgroup and 10 HCs in the balanced subgroup from Ext-DS, using the Brunner-Munzel test.

## Results

### Identification of the OCD Network

To explore the neural basis of OCD and related imbalanced conditions (*ν*^+^ > *ν*^−^ ), we examined whole-brain FC using rs-fMRI. We hypothesized that a shared neural signature, present in non-medicated OCD patients and extending to a subgroup of HCs with imbalanced learning, would be detected.

First, a whole-brain FC matrix using the CAB-NP was constructed and compared between 49 non-medicated OCD patients and 53 HCs (Core-DS; see Methods and Supplementary Table 1) using NBS. This analysis, with an initial threshold of t = 3.87 and 10,000 random permutations (see Methods), detected a significant network component related to OCD (the ‘OCD network’), exhibiting significantly increased connectivity in non-medicated OCD patients compared with HCs (p_adjusted_ = 0.022, Figure 1). The OCD network comprised nodes in the DLPFC, parietal cortex, RSC, and the hippocampal formation and parahippocampal region. These regions are primarily associated with the default mode network (DMN) and the frontoparietal network (FPN), as detailed in Supplementary Figure 1.

**Figure 1.**
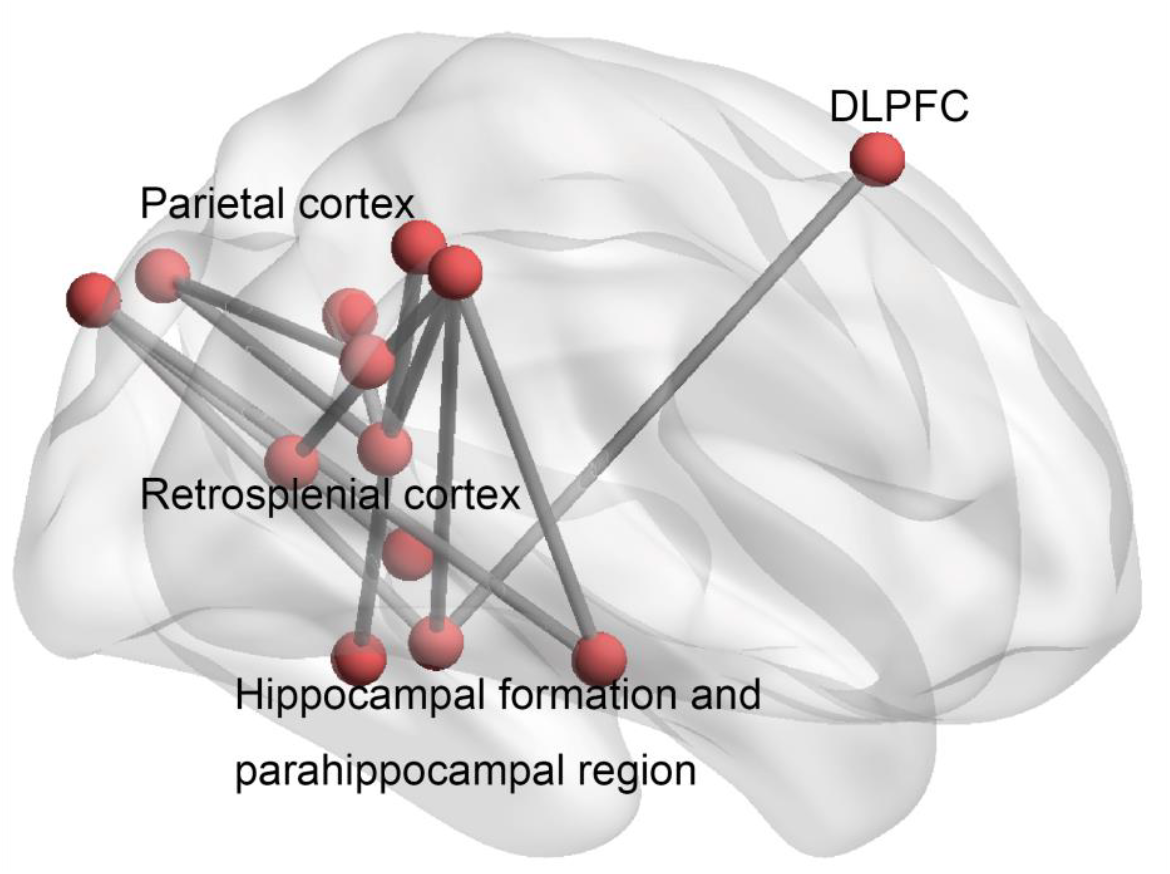
The identified OCD network. A significantly increased functional network component in non-medicated OCD participants, compared with HCs (p_adjusted_ = 0.022).

### Validation of the OCD Network

To validate the identified OCD network, mean FC in this network was compared between non-medicated OCD patients (n = 10) and HCs (n = 18) in the IndV-DS (Supplementary Table 2). Consistent with initial findings, mean FC in the ‘OCD network’ was significantly greater in non-medicated OCD patients than in HCs (Figure 2, Brunner-Munzel test, statistic = -3.11, p = 0.0027).

**Figure 2.**
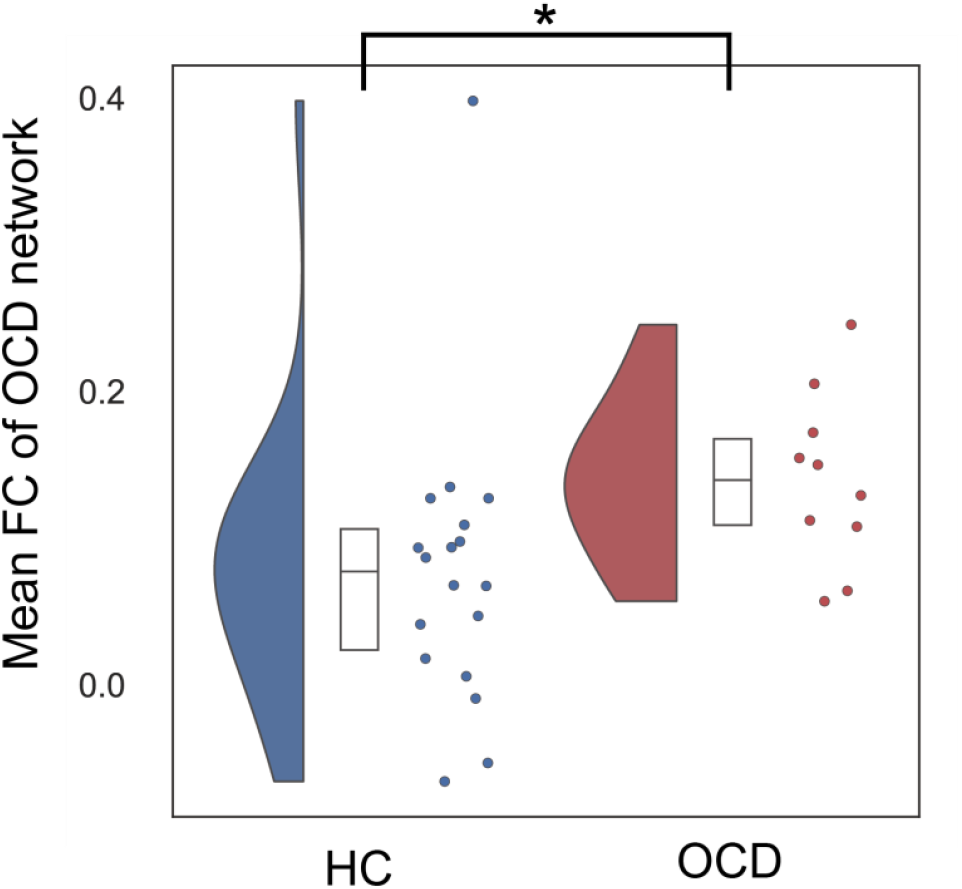
The Validation of the OCD Network. Significantly increased mean FC in the OCD network of non-medicated OCD patients compared with HCs in the entirely independent dataset (* Brunner-Munzel test, statistic = −3.11, p = 0.0027). The half-violin, box, and strip plots represent the probability density, interquartile range and median, and raw data, respectively.

### Neural Correlates of Imbalanced Learning in HCs

Finally, we investigated whether the imbalanced (*ν*^+^ > *ν*^−^) HC subgroup also showed OCD-like FC patterns. All FCs of the OCD network were compared between 10 HCs in the imbalanced cluster and 10 HCs in the balanced cluster (Ext-DS, independent of Core-DS and IndV-DS; see Methods and Supplementary Table 3), previously identified by our delayed feedback task ^6^. We found that FC between the DLPFC and presubiculum was significantly increased in the imbalanced HC subgroup, similar to the OCD group (Figure 3, Brunner-Munzel test, statistic = -2.94, p = 0.0051). A trend for increased connectivity was maintained after correcting multiple comparisons (p_adjusted_ = 0.066).

**Figure 3.**
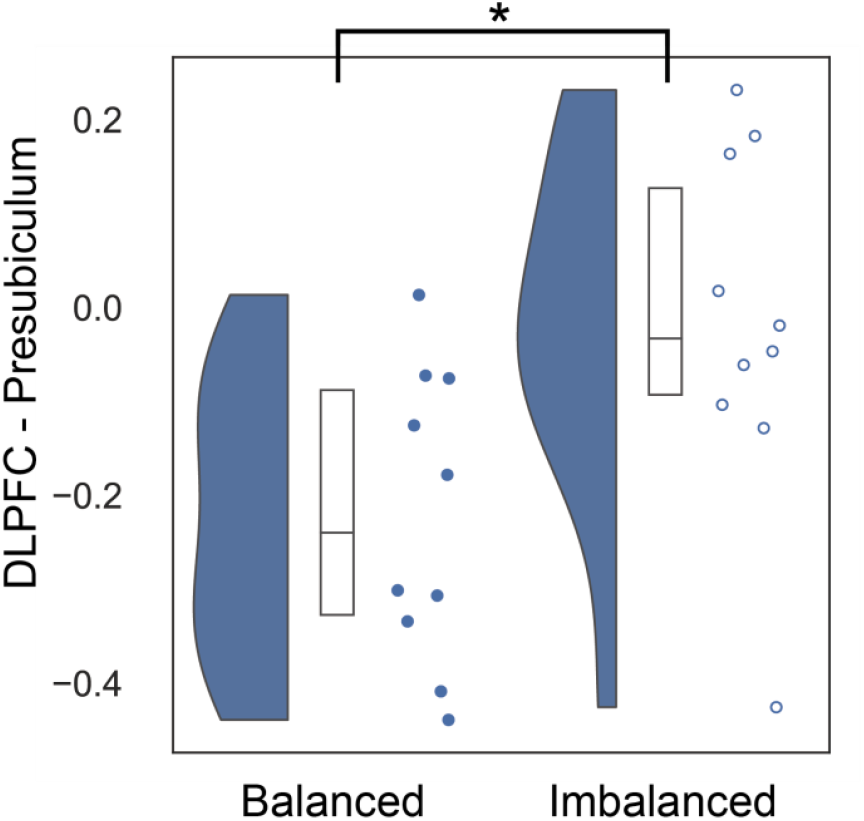
Neural Correlates of Imbalanced Learning in HCs. The imbalanced HC group showed OCD-like increased FC between the DLPFC and presubiculum (* Brunner-Munzel test, statistic = −2.94, p = 0.0051). The half-violin, box, and strip plots represent the probability density, interquartile range and median, and raw data, respectively.

## Discussion

This study revealed two key findings: first, we identified the ‘OCD network,’ showing increased FC in OCD patients; and second, we found increased connectivity in this network, particularly between the DLPFC and presubiculum, in a subgroup of healthy individuals with imbalanced learning. These results provide new insights into neural substrates of OCD and how they relate to the behavioral dimension of imbalanced reinforcement learning.

Identification of the ‘OCD network,’ exhibiting increased FC in OCD patients, is a significant finding, particularly given its replication in an independent dataset. Using NBS, we found that the ‘OCD network’ consisted of regions in the DLPFC, parietal cortex, RSC, and hippocampal formation. Specifically, the ‘OCD network’ showed increased FCs in the DMN (7 of 13) and between the DMN and FPN (4 of 13) (Supplementary Figure 1). Although traditional theories of OCD often focus on the cortico-striato-thalamo-cortical (CSTC) circuit ^15^, our findings, consistent with our previous work ^3^, highlight the importance of the DMN and FPN. These regions, as also discussed in Gürsel et al.’s meta-analysis ^5^, which demonstrates the importance of frontoparietal regions’ intrinsic connectivity in OCD, may be critical in OCD pathophysiology by impacting a wider range of self-referential thought processes. Furthermore, our previous work showed that imbalanced trace factors are crucial for OCD mechanisms, and this identified widespread altered network may represent a critical neural substrate for this imbalance ^6^.

Extending beyond OCD, our findings also revealed increased FC between the DLPFC and presubiculum in the OCD network in a subgroup of healthy individuals exhibiting imbalanced learning (*ν*^+^ > *ν*^−^), similar to that seen in OCD patients. This finding highlights a continuous neural characteristic, linking a behaviorally defined learning imbalance to altered FC. While the increase in FC did not survive correction for multiple comparisons across all connections in the OCD network, the strong trend (p_adjusted_ = 0.066) is still compelling. The DLPFC, a key component of the OCD network, is implicated in processing information relevant for credit assignment ^11^ and for forming associations between items during online processing ^26^. Similarly, the RSC, also part of the identified network and interconnected with the hippocampus, is thought to be important for memory retrieval and its influence on behavior ^13^. Considering the roles of the DLPFC, RSC, and hippocampus in reinforcement learning, increased connectivity between the DLPFC and presubiculum in the imbalanced HC subgroup may represent a critical neural substrate for the observed learning imbalance. Our results suggest that altered connectivity in the ‘OCD network’ may contribute to core neural processes underlying both OCD and subclinical obsessive-compulsive traits.

These findings have several important implications ^27^. First, the identification of a continuous neural substrate for imbalanced learning in both OCD patients and a subgroup of HCs suggests that OCD traits exist on a spectrum rather than as a purely discrete pathological condition, supporting the idea of a broader spectrum for mental disorders ^28^. Furthermore, combining computational modeling and whole-brain FC data constitutes a powerful method for dissecting complex psychiatric disorders. Moreover, the identification of a continuous network in healthy individuals with imbalanced memory traces may represent a potential endophenotype for OCD, where similar neural characteristics are present in individuals regardless of a clinical diagnosis. Further studies using this approach should evaluate specific network features found in our research in unaffected family members, which may help reveal genetic underpinnings and neural mechanisms of OCD ^27, 29-31^.

In summary, this study identified a key functional network, the ‘OCD network,’ showing altered connectivity in OCD and significantly demonstrated that this network also participates in a subgroup of HCs with behaviorally defined learning imbalances. Our data-driven approach, which combined resting-state fMRI and reinforcement learning-based parameterization of behavior, provides a novel way to bridge computational and neural levels of analysis in mental disorders. However, this study had several limitations. Finally, although the model demonstrated the importance of identified regions in the brain network, future studies are needed to examine this in a longitudinal framework. Future research should also investigate other factors that influence the trajectory of compulsivity, such as environmental influences. Further research using larger samples, longitudinal study designs, and interventional approaches could give more insight into the nature and treatment of OCD and other conditions on the obsessive-compulsive spectrum.

## Conclusion

This study investigated neural underpinnings of imbalanced learning, a phenomenon implicated in OCD, using resting-state FC. We identified the ‘OCD network’ with altered connectivity in OCD patients and, importantly, found that increased connectivity in this network, especially between the DLPFC and presubiculum, was also present in a subgroup of healthy individuals exhibiting imbalanced learning. These results suggest that neural correlates of OCD can be observed among both OCD patients and a subgroup of healthy individuals. These findings support the model that this network represents a shared neural basis for behavioral manifestations of imbalanced reinforcement learning connected to the obsessive-compulsive spectrum. Our data-driven approach, combining resting-state fMRI with a reinforcement learning framework, reveals a potentially important link between a computational model and large-scale brain networks. Future research should examine causal relationships between these network alterations and learning imbalances and should assess potential manipulation of these mechanisms to develop treatment and prevention strategies for OCD and other related conditions. Longitudinal studies and interventions targeting these networks and the influence of environmental factors are warranted to gain a comprehensive understanding of the dynamics of compulsivity.

## Data Availability

Patient data supporting the conclusions of this paper are not publicly available because they contain information that could compromise participant privacy or consent.

## Acknowledgments

All authors had full access to the data used in this study, and the corresponding author, S.C. Tanaka, takes responsibility for data integrity and accuracy of the data analysis. These research results have been achieved by JSPS KAKENHI Grant Number JP24H00069 (S.C. Tanaka). The following partly supported this study: “Research and development of technology for enhancing functional recovery of elderly and disabled people based on non-invasive brain imaging and robotic assistive devices”, the Commissioned Research of National Institute of Information and Communications Technology (NICT), JAPAN (S.C. Tanaka); AMED under Grant Number JP23wm0625001 (S.C. Tanaka); JSPS KAKENHI Grant Number JP16K01958, JP16H06396, and JP21H05172 (S.C. Tanaka); JSPS KAKENHI Grant Number JP16H01516, JP18H05524, and JP23H00494 (Yutaka Sakai); JSPS KAKENHI Grant Number 16H01512 and the Sakamoto Research Foundation of Psychiatric Diseases (Yuki Sakai); the Joint Usage/Research Center (“Behavioral economics”) of Institute of Social and Economic Research, Osaka University (J. Narumoto).

## Disclosure statement

The authors declare no competing financial interests.

## Author Contributions

Yuki S, Yutaka S, JN, and SCT designed the study. The theoretical framework was based on the idea developed by Yutaka S and Yuki S. Yuki S, YA, JN, and SCT collected the data. Yuki S conducted computational modeling and statistical analysis of the data. Yuki S wrote the manuscript, which all authors edited. SCT acquired funding to support theory development and data analysis.

## Notes

### Competing Interest Statement

The authors have declared no competing interest.

