## Supplementary information for "“Continuous Neural Correlates of Imbalanced Reinforcement Learning in Obsessive-Compulsive Disorder and Healthy Individuals”"

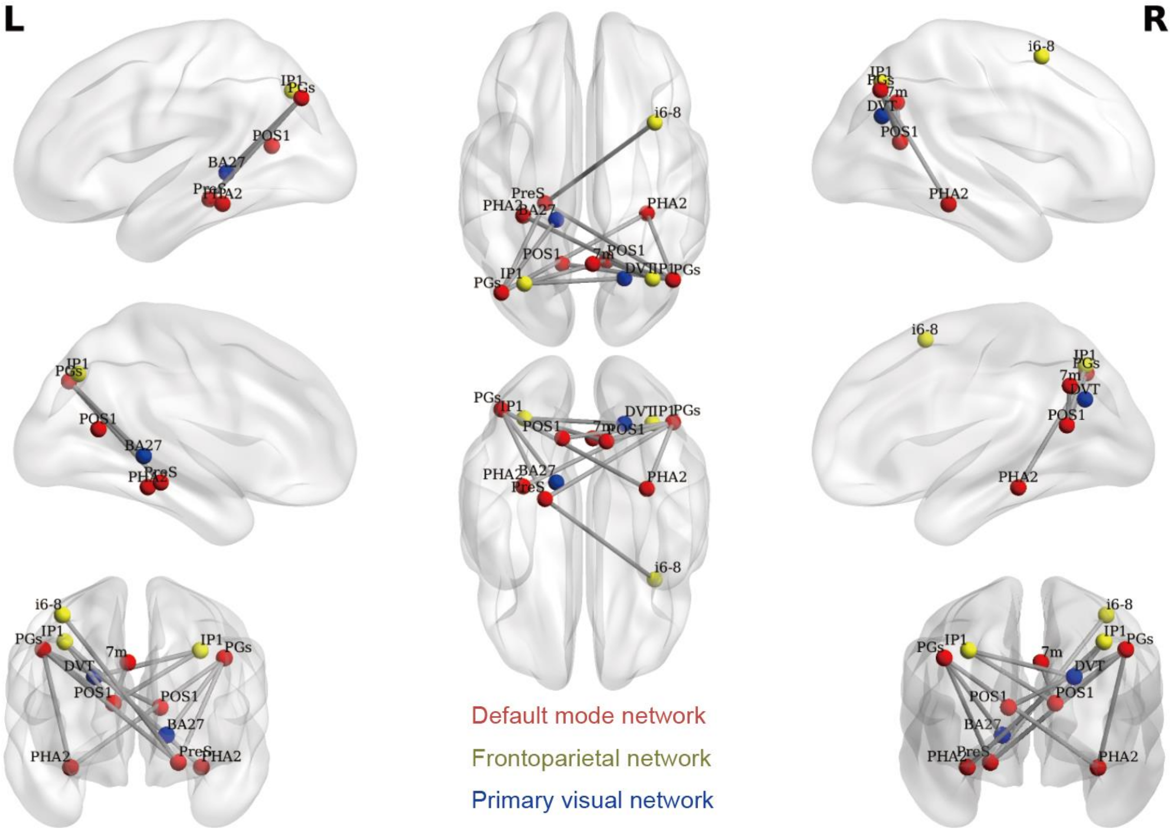


**Supplementary Figure 1. Detailed view of the OCD network.** L and R represent left and right, respectively. The network color of each node corresponds to that of CAB-NP[1]. Abbreviations of nodes are based mainly on parcellation developed by Glasser et al.[2] [Human Connectome Project Multi-Modal Parcellation (HCP-MMP)], except for one node (BA27) that is only included in CAB-NP. All functional connections are listed below in the form of "Area Name (Area Description)" based on the supplementary information of HCP-MMP [2]. The area name of BA27 (Brodmann area 27) was based on the Montreal Neurological Institute coordinate of the CAB-NP parcel.

1. L, IP1 (Area IntraParietal 1) – R, POS1 (Parieto-Occipital Sulcus Area 1)
2. R, i6-8 (Inferior 6–8 Transitional Area) – L, Pres (PreSubiculum)

3. R, IP1 (Area IntraParietal 1) – L, PHA2 (ParaHippocampal Area 2)
4. R, IP1 (Area IntraParietal 1) – R, POS1 (Parieto-Occipital Sulcus Area 1)
5. L, POS1 (Parieto-Occipital Sulcus Area 1) – R, PGs (Area PGs)
6. L, Pres (PreSubiculum) − L, PGs (Area PGs)
7. L, Pres (PreSubiculum) − R, PGs (Area PGs)
8. L, PGs (Area PGs) – R, PHA2 (ParaHippocampal Area 2)
9. R, 7m (Area 7m) − R, POS1 (Parieto-Occipital Sulcus Area 1)
10. R, POS1 (Parieto-Occipital Sulcus Area 1) − R, PGs (Area PGs)
11. R, PGs (Area PGs) − R, PHA2 (ParaHippocampal Area 2)
12. R, DVT (Dorsal Transitional Visual Area) – L, IP1 (Area IntraParietal 1)
13. L, BA27 (Brodmann area 27) – L, PGs (Area PGs)

**Supplementary Table 1. Demographic information of Core-DS.**

All demographic distributions are matched between the OCD and HC populations ($p > 0.05$).

| | **OCD patients (n = 49)** | **Healthy controls (n = 53)** |
|---|---|---|
| **Demographic characteristics** | | |
| Age, years | 32.5 ± 9.9 | 29.4 ± 7.4 |
| Male sex, No. | 20 | 26 |
| Left-handed†, No. | 3 | 2 |
| **Clinical characteristics** | | |
| With/without medication, No. | 0 / 49 | NA |
| Total Y-BOCS score† | 22.1 ± 6.2 | NR |

Values represent the mean ± standard deviation.

Abbreviations: Core-DS, Core Discovery Dataset; NA, not applicable; NR, not recorded; OCD, obsessive-compulsive disorder; Y-BOCS, Yale-Brown Obsessive-Compulsive Scale[3].

†Classification of handedness was based on a modified 25-item version of the Edinburgh Handedness Inventory. All patients were surveyed for obsessive-compulsive symptoms using the Japanese version of Y-BOCS.

**Supplementary Table 2. Demographic information of IndV-DS.**

All demographic distributions are matched between the OCD and HC populations ($p > 0.05$).

| | **OCD patients (n = 10)** | **Healthy controls (n = 18)** |
|---|---|---|
| **Demographic characteristics** | | |
| Age, years | 31.6 ± 10.4 | 29.9 ± 8.7 |
| Male sex, No. | 4 | 8 |
| Left-handed†, No. | 1 | 3 |
| **Clinical characteristics** | | |
| With/without medication, No. | 0 / 10 | NA |
| Total Y-BOCS score† | 23.8 ± 5.8 | NR |

Values represent the mean ± standard deviation.

Abbreviations: IndV-DS, Independent Validation Dataset; NA, not applicable; NR, not recorded; OCD, obsessive-compulsive disorder; Y-BOCS, Yale-Brown Obsessive-Compulsive Scale[3].

†Classification of handedness was based on a modified 25-item version of the Edinburgh Handedness Inventory. All patients were surveyed for obsessive-compulsive symptoms by the Japanese version of Y-BOCS.

**Supplementary Table 3. Demographic information of the rs-fMRI Ext-DS.**

All demographic distributions are matched between the unbalanced and balanced clusters ($p > 0.05$).

| | **Imbalanced group (n = 10)** | **Balanced group (n = 10)** |
|---|---|---|
| **Demographic characteristics** | | |
| Age, years | 22.8 ± 2.1 | 22.0 ± 1.5 |
| Male sex, No. | 9 | 9 |
| Left-handed†, No. | 2 | 0 |

Values represent the mean ± standard deviation.

Abbreviations: Ext-DS, Extension Dataset.

†Classification of handedness was based on a modified 25-item version of the Edinburgh Handedness Inventory.

**Supplementary Table 4. Summary of imaging protocols for resting-state fMRI at the three sites.**

| Parameter | Core-DS (Kajiicho Medical Imaging Center) | IndV-DS (Kyoto Prefectural University of Medicine) | Ext-DS (ATR) |
|---|---|---|---|
| MRI scanner | Philips Achieva | Gyroscan Intera | Siemens Verio |
| Magnetic field strength, T | 3 | 1.5 | 3 |
| Field of view, mm | 192 | 192 | 212 |
| Matrix | 64 × 64 | 64 × 64 | 64 × 64 |
| Number of slices | 39 | 35 | 39 |
| Number of volumes | 200 | 200 | 244 |
| Slice thickness, mm | 3 | 3.6 | 3.2 |
| Slice gap, mm | 0 | 0 | 0.8 |
| TR, ms | 2000 | 2411 | 2500 |
| TE, ms | 30 | 40 | 30 |
| Flip angle, ° | 80 | 80 | 80 |

| | | | |
|---|---|---|---|
| Instruction to participants and other imaging conditions | Participants were instructed simply to keep their eyes closed, not to think of anything in particular, and not to fall asleep. | Participants were instructed simply to keep their eyes closed, not to think of anything in particular, and not to fall asleep. | Participants were instructed simply to keep looking at the crosshair mark presented, not to think of anything in particular, and not to fall asleep. |
